## Supplementary Materials for "Analytical validity of nanopore sequencing for rapid SARS-CoV-2 genome analysis"

**Fig. S1. Performance metrics for ONT and Illumina sequencing of synthetic SARS-CoV-2 controls.** p.2

**Fig. S2. Erroneous variants detected in ONT sequencing of synthetic SARS-CoV-2 controls.** p.3

**Fig. S3. False-positive and false-negative variants at error-prone low-complexity sequence.** p.4

**Fig. S4. Examples of large deletions detected by ONT sequencing of SARS-CoV-2 specimens.** p.5

**Supplementary Table 1.** Variant 'truth' sets generated by Illumina sequencing on 157 SARS-CoV-2 specimens**.**

Table provided as separate Excel file.

**Supplementary Table 2.** Detection of consensus-level variation in individual SARS-CoV-2 specimens with ONT sequencing.

Table provided as separate Excel file.

**Supplementary Table 3.** Impact of excluding low-complexity 'blacklist' sites on consensus variant detection in individual SARS-CoV-2 specimens.

Table provided as separate Excel file.

**Supplementary Table 4.** Pairwise comparisons of technical replicates for Illumina and ONT SARS-CoV-2 WGS.

Table provided as separate Excel file.

**Supplementary Table 5.** Detection of sub-consensus variation in individual SARS-CoV-2 specimens with ONT sequencing.

Table provided as separate Excel file.

**Supplementary Table 6.** List of sequencing runs performed in the study.

Table provided as separate Excel file.

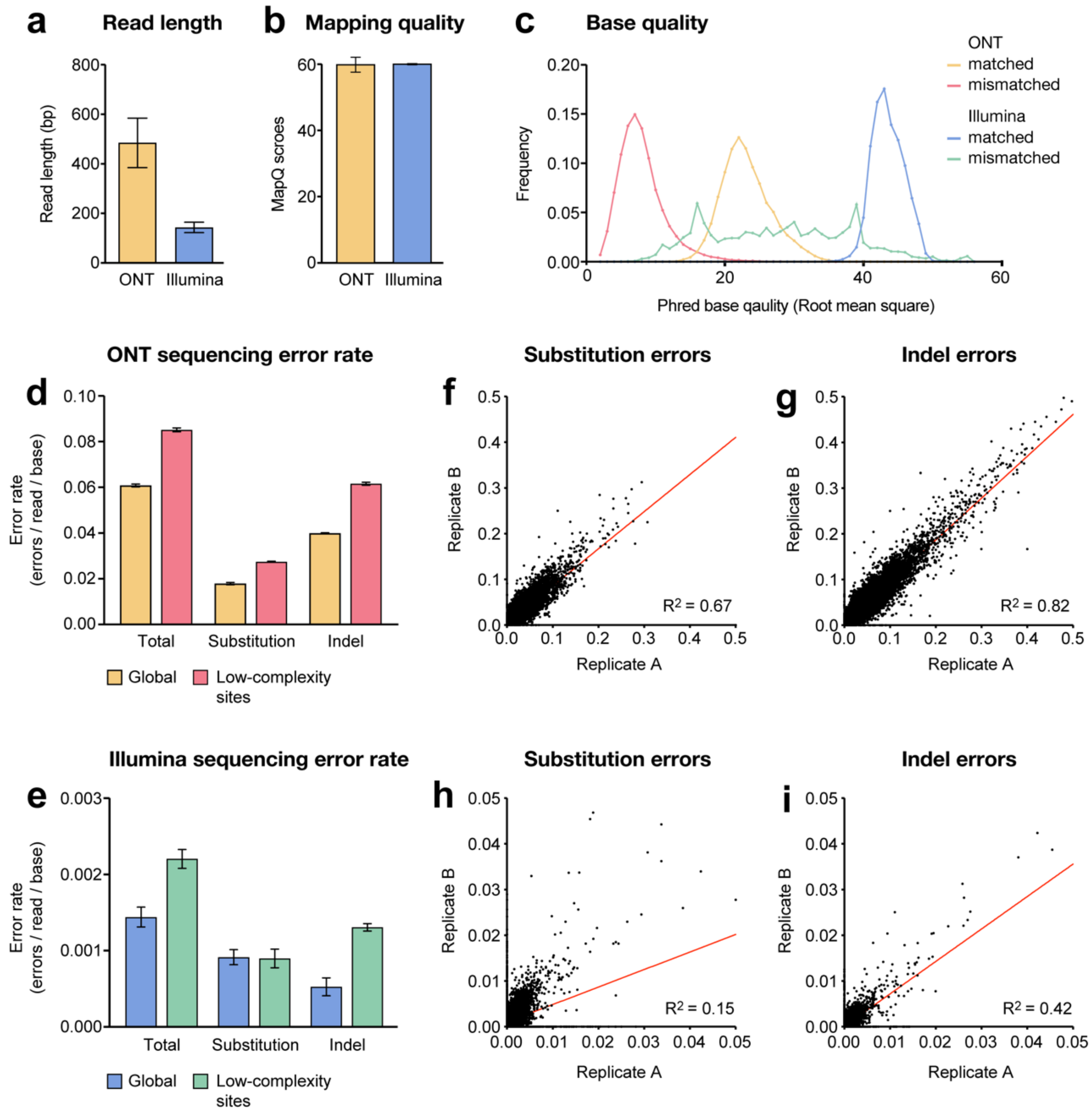


**Fig. S1. Performance metrics for ONT and Illumina sequencing of synthetic SARS-CoV-2 controls.** Read lengths (**a**) and MapQ scores (**b**) for ONT (gold) and Illumina (blue) sequencing reads. Bars show mean ± 95% CI. (**c**) Frequency distributions show Phred base quality scores (Root mean square) within ONT and Illumina sequencing alignments. Bases that were mismatched to the reference sequence, indicative of sequencing errors, are shown separately from bases that matched the reference. (**d,e**) Read-level error rates for ONT (**d**) and Illumina (**e**) sequencing reads. Error rates are shown separately for the complete SARS-CoV-2 genome, and low-complexity sites (*n* = 15, approx. 1% of the genome) where elevated error rates were observed. Bars show mean ± standard deviation across 8 replicates for each technology. (**f-i**) Scatter plot show correlation in per-base error frequency profiles between replicates for ONT (**f,g**) and Illumina (**h,i**). Substitution and indel errors plotted separately. Correlation in error frequency profiles between replicates indicates profiles are non-random, with local sequence context influencing error rates.

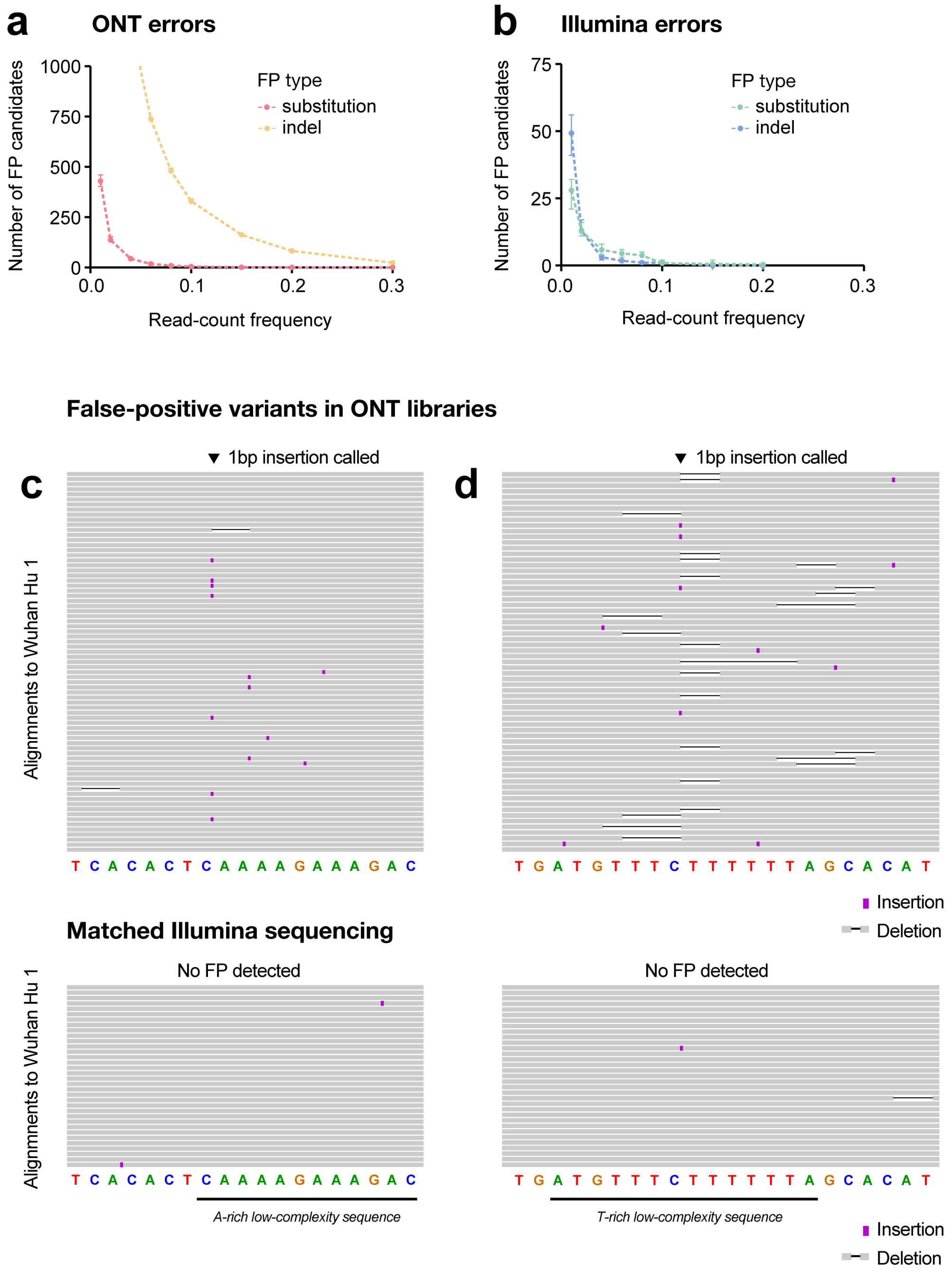


**Fig. S2. Erroneous variants detected in ONT sequencing of synthetic SARS-CoV-2 controls.** (**a,b**) Curves show the number of substitution and indel errors detected in synthetic RNA controls relative to their read count frequencies. (**c,d**) Genome browser views of two single-base insertions that were erroneously detected by ONT sequencing (upper panels) on with SARS-CoV-2 controls. Both erroneous insertions were called at error-prone low-complexity sequences and neither was supported by Illumina sequencing alignments (lower panels) at the same regions.

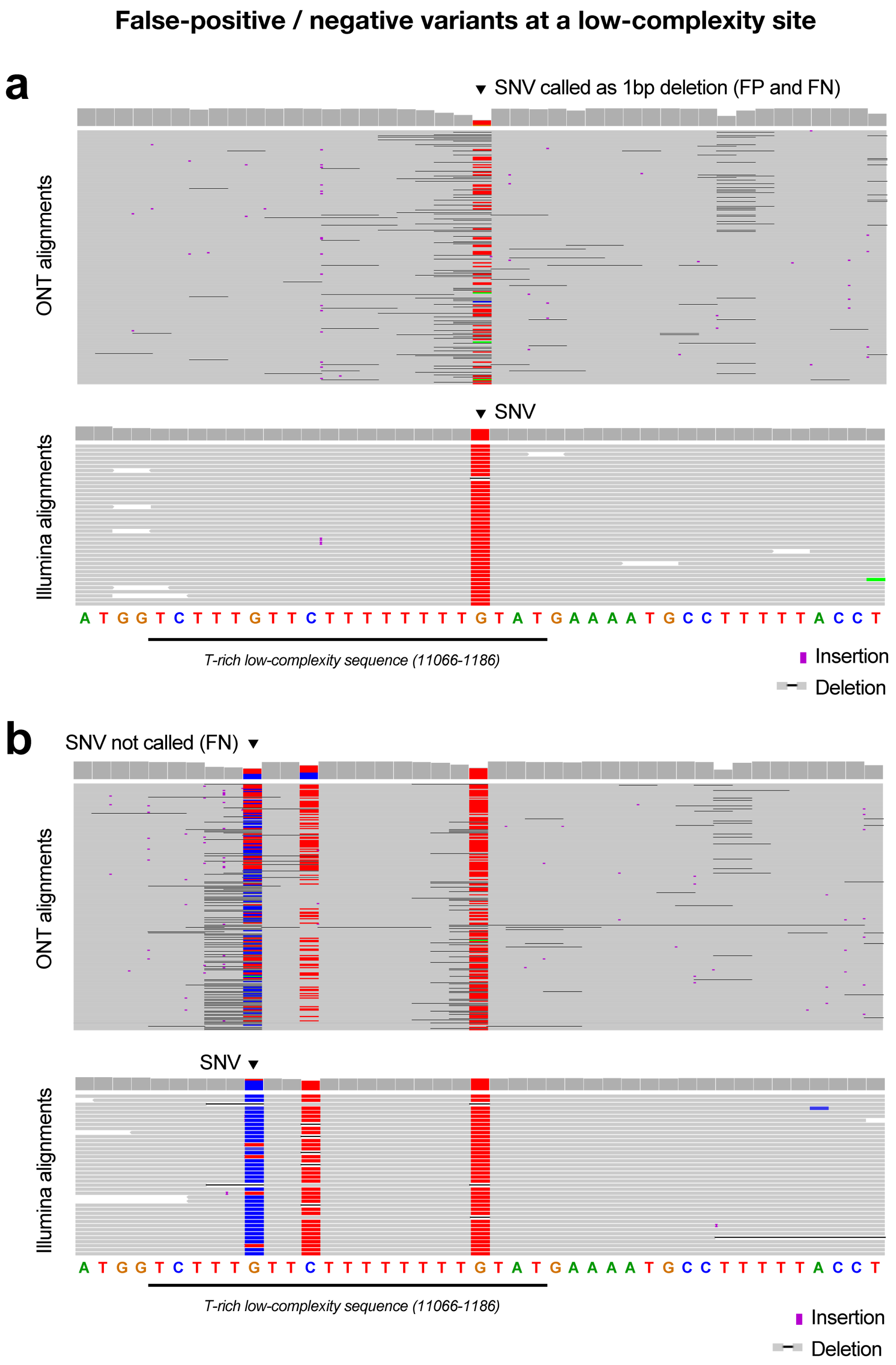


**Fig. S3. False-positive and false-negative variants at error-prone low-complexity sequence.** (**a**,**b**) Genome browser views show ONT (upper) and Illumina (lower) sequencing alignments at a T-rich low-complexity site in the *orf1ab* gene, where false-positives and false-negatives were found in multiple specimens. In example **a**, an SNV identified by Illumina reads was erroneously interpreted as a 1 bp deletion. In example **b**, the leftmost SNV was not detected by ONT due to confounding errors at the same position.

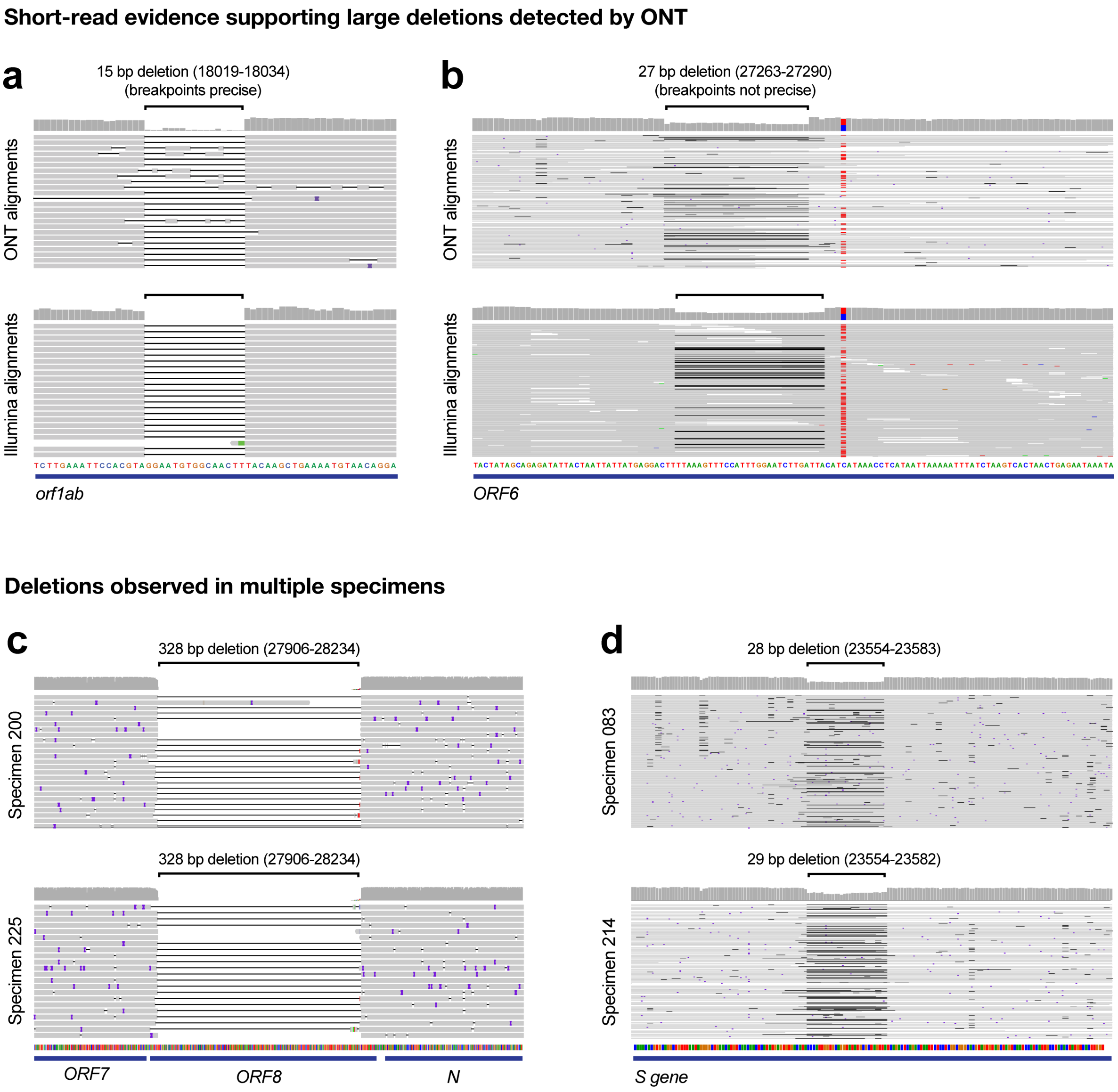


**Fig. S4. Examples of large deletions detected by ONT sequencing of SARS-CoV-2 specimens.** (**a,b**) Genome browser views show ONT (upper) and Illumina (lower) sequencing alignments at the site of deletions detected in *orf1ab* (**a**) and *ORF6* (**b**). In both cases, evidence for split short-read alignments support the candidate variant detected by ONT sequencing. For the deletion in **a**, breakpoints were accurately resolved by ONT sequencing, whereas the breakpoints identified in **b** are not precise. (**c**) Genome browser views shows 2 identical 328 bp deletions in *ORF8* that were identified in multiple independent specimens. (**d**) Genome browser views shows 2 highly similar, but non-identical, 28 and 29 bp deletions in *S* that were identified in multiple independent specimens.
